# More pests but less treatments: ambivalent effect of landscape complexity on Conservation Biological Control

**DOI:** 10.1101/2021.03.19.436155

**Authors:** Patrizia Zamberletti, Khadija Sabir, Thomas Opitz, Olivier Bonnefon, Edith Gabriel, Julien Papaïx

## Abstract

In agricultural landscapes, the amount and organization of crops and semi-natural habitats (SNH) have the potential to promote a bundle of ecosystem services due to their influence on ecological community at multiple spatio-temporal scales. SNH are relatively undisturbed and are often source of complementary resources and refuges, supporting more diverse and abundant natural pest enemies. However, the nexus of SNH proportion and organization with pest suppression is not trivial. It is thus crucial to understand how the behavior of pest and auxiliary species, the underlying landscape structure, and their interaction may influence conservation biological control (CBC). Here, we develop a generative stochastic landscape model to simulate realistic agricultural landscape compositions and configurations of fields and linear elements. Generated landscapes are used as spatial support over which we simulate a spatially explicit predator-prey dynamic model. We find that SNH boost predator population, but predator movement from hedges to fields is fundamental for an efficient pest regulation by auxiliaries and to decrease pesticide treatments. Moreover landscape elements may lead to different effects on pest reduction depending on the considered scale. Integration of species behaviors and traits with landscape structure at multiple scales are needed to provide useful insights for CBC.

## Introduction

Agricultural landscape simplification results in substantial loss of semi-natural mosaics and of non-crop field margins. It is often associated with high pest abundance, which in turn requires a higher pesticide input (1,2). Consequently, a negative relationship emerges between intensity of agriculture and agricultural landscape biodiversity (3) because of a partial replacement and suppression of the ecological services provided by communities of beneficial organisms (4,5). Habitat heterogeneity is key to allow cross-system fluxes of organisms across agro-ecological interfaces by influencing ecological dynamics within those habitats (6,7) and potentially increasing natural enemy abundance and diversity in agricultural systems (8,9). In addition, complex landscape favours habitat and resource diversity for natural enemies with increased availability of alternative preys, higher microclimate heterogeneity, the presence of refuges from their own predators and for overwintering (10). In arable land, semi-natural habitat (SNH) is typically restricted to hedgerows. These linear structures play an important role as relatively perennial line corridors because of their temporal stability with respect to crop fields. Their presence supports natural enemy dispersal and movement to escape from disturbances and to find food resources scattered in time and space (11,12).

While SNH favours the presence or abundance of functional groups of organisms in landscapes, it can also results in ineffective conservation biological control (CBC) (12,13) with no, or even negative effects on pest control (12–14). A meta-analysis revealed that pest pressure in complex landscapes is reduced in 45% of cases, not affected in 40% of cases and increased in 15% of cases (9). For example, the effect of landscape structure on pests themselves remains inconclusive, as many crop pests also benefit from nearby non-crop habitat (12–14). In particular, SNH can offer more complementary resources to pests rather than to natural enemies to complete their life cycle (6). Predator abundance is not always enough to guarantee a consistent reduction of pest species (15) in case of the presence of alternative prey (known as *dilution effect*) (16), or increased intra-guild predation (17). Life history traits, in particular those traits related to mating systems, competitive skills, movement abilities and habitat use, are also of major importance by affecting species responses to landscape heterogeneity and being readily linked with ecological processes (18). Thus, effect direction and magnitude jointly depend on organisms and landscapes under study (19,20).

The impacts of landscape structure on pest population dynamics are generally investigated through empirical correlative approaches with global descriptors at landscape level, due to the difficulty of manipulating large landscapes for local analyses and for the lack of the spatio-temporal dimension. The main drawback of these approaches is the difficulty of linking correlation levels to population dynamic processes such as local population growth or migration behaviour (21). A complementary approach consists in coupling generative landscape models with population dynamics models to explore how different landscape configurations, including the hedge network structure, affect CBC (22).

In this work, we develop a stochastic model to generate realistic landscapes, where we simulate a spatially explicit population dynamic model of a predator-pest system integrating dispersal both on agricultural fields and on hedge network. A major goal is to implement a general simulation-based approach to obtain theoretical insights on CBC by incorporating landscape effects and species traits, which can serve as basis to formulate practical recommendations. Specifically, following questions are investigated: (i) What are the main factors that influence the predator-pest population densities in complex landscape? (ii) How do life history traits modify the effect of landscape heterogeneity? (iii) Can landscape heterogeneity reduce number of pesticide applications by enhancing CBC?

## 2 Models and methods

### 2.1 Stochastic landscape model

The landscape is represented through a vectorial approach, which is appropriate for representing the highly regular geometrical patterns of agricultural landscapes (23,24). It is composed of polygons representing fields, separated by edges. Landscape elements are characterized by their geometry (e.g., vertex coordinates, size and shape), and by categorical information defining the land-cover (e.g., crop or natural habitat). The landscape geometrical structure is fixed and based on a real landscape with an extent of 5.55 km. The landscape is transformed into a T-tessellation (25,26) composed of 188 polygons with a total of 577 edges.

We use Gaussian random fields (GRFs) to allocate a proportion of polygons and edges as crops representing the principal culture and hedges to provide SNHs, respectively. A threshold on the simulated GRF values is set to attribute specific landscape elements depending on the value being below or above the threshold. By definition, a GRF denoted by *W* is a random surface over continuous 2D space, for which the multivariate distribution of the values (*W*(*x*_1_),*W*(*x*_2_),…,*W*(*x*_*n*_)) observed at a finite number of locations *x*_1_,*x*_2_,…,*x*_*n*_ in the landscape corresponds to a multivariate normal distribution, characterized by its mean vector and its covariance matrix Σ. The mean is fixed to 0 and the exponential correlation function is used for Σ, such as 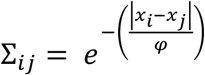, where |*x*_*j*_ − *x*_*i*_| is the Euclidean distance between any two points *x*_*j*_ and *x*_*i*_. The range parameter *φ* ≥ 0 governs the strength of clustering of category allocation to landscape elements. To handle the interactions between the allocation of hedge and crop, we simulated two correlated GRFs for crop (*W*_*c*_(*s*)) and hedge (*W*_*h*_(*s*)):

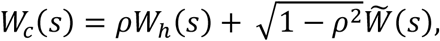

where *ρ* ∈ [−1,1] controls the correlation between *W*_*h*_ and 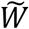, which is a GRF independent from *W*_*h*_. The parameter used for the landscape models with their range of values can be found in Table 1. Figure 1 shows an example of four landscapes simulated according to different proportions and aggregation levels of hedges and crop fields.

**Table 1:**
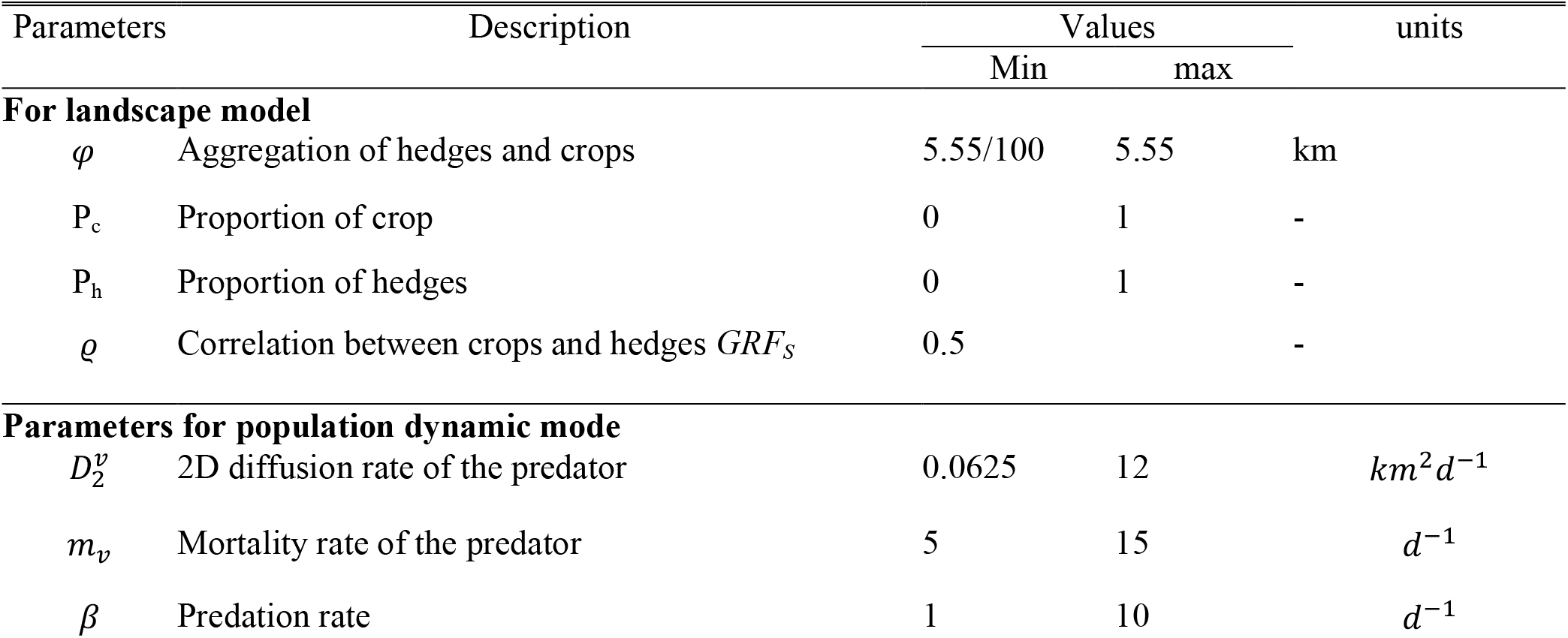
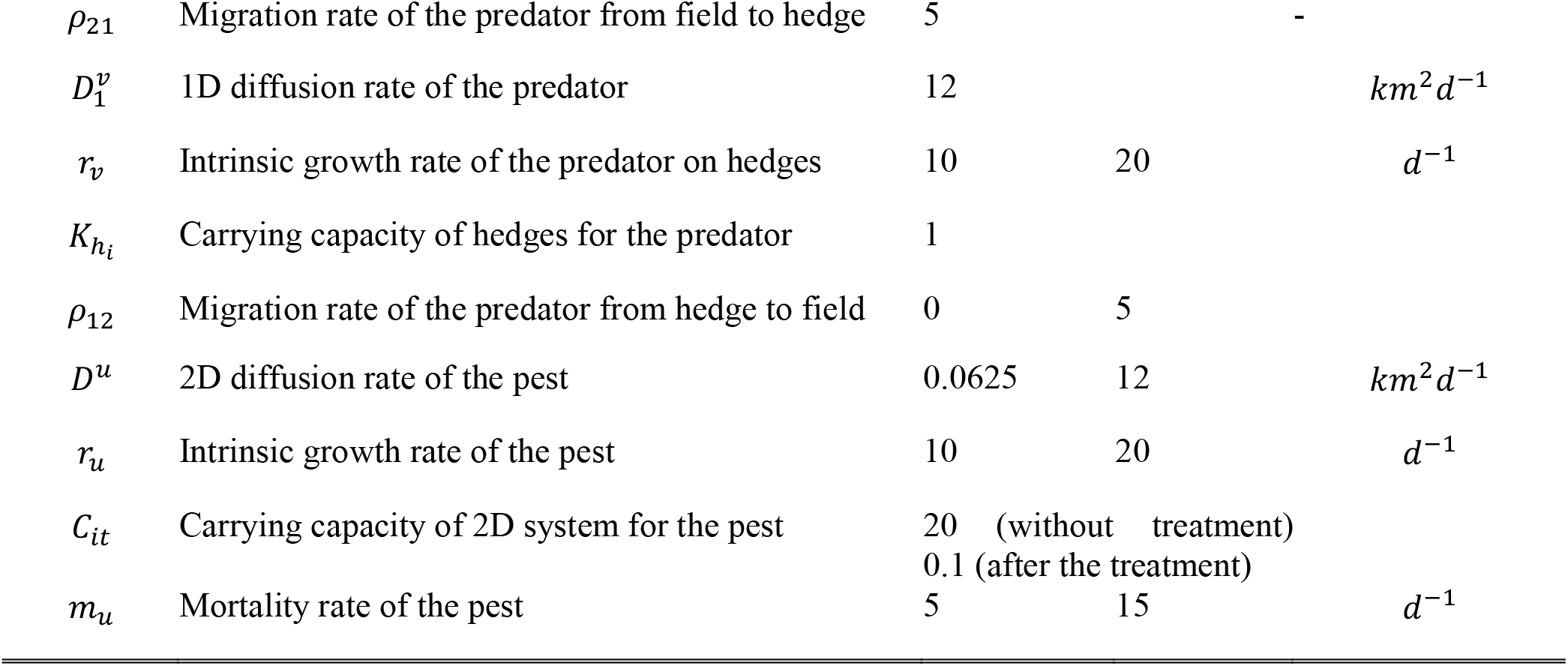
Description of parameter values.

**Figure 1:**
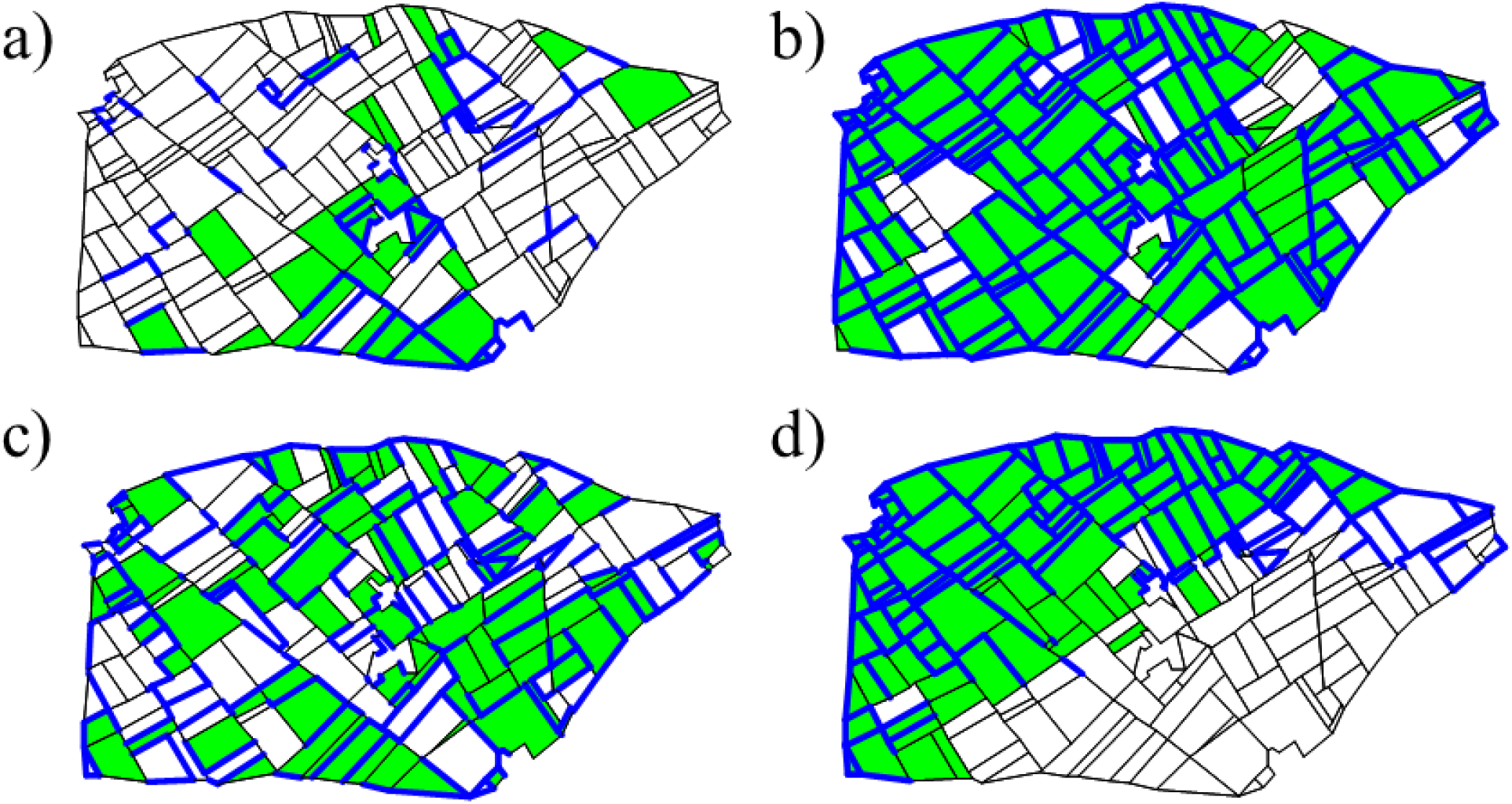
Examples of simulated landscape structures with interacting elements using the following allocation categories: for fields, (i) crop (green) and (ii) non-crop (white); for edges, (i) hedge (blue) and (ii) no-hedge (black). First row: low (a) and high (b) proportions of crop and hedges (0.2 and 0.8, respectively), with fixed parameter configuration for aggregation and fixed correlation between crop and hedges (0.5). Second row: low (c) and high (d) crop and hedge aggregation level from left to right, with fixed proportion of crop and hedges (0.5) and fixed correlation between crop and hedges (0.5).

### 2.2 Predator-pest model

We developed a spatially explicit predator-pest model based on a system of partial differential equations. The model is built on a previously developed model that considers both 2D diffusion on polygons and 1D diffusion on edges (11). Simulations are performed over a [0,1]-time interval representing a cropping season of several months with a time step of 0.01, such that the time unit corresponds to one day. The model parameters and their range of simulated values are reported in Table 1. Numerical simulations of the spatio-temporal partial differential equation system of pest-predator dynamics are performed using the Freefem++ finite-element framework (27).

### Predator dynamics

Hedges are the main habitat of the predator. Using notations *t* for time and *x* for a spatial location, we thus assume the following 1-dimensional reaction-diffusion model for the predator density 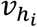 on the edge *h*_*i*_:

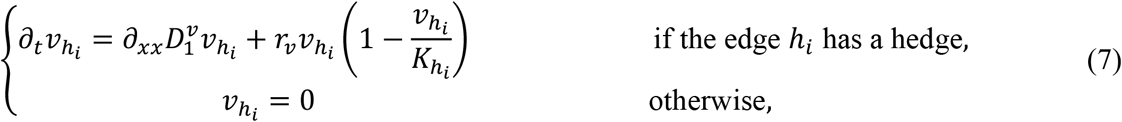

Where 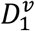 is the diffusion parameter of the predator along hedges, *r*_*ν*_ is the intrinsic growth rate of the predator, and 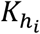 is the carrying capacity of the hedge *i*. If two hedges are linked together at one of their endpoints, then the dynamics in Equation (7) apply continuously across the junction.

In addition, the predator forages on fields where it feeds on the pest. The population density 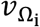 of predators in a field Ω_*i*_ is modelled by a reaction-diffusion equation with mobility parameter within field 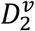, predation rate *β*, and mortality *m*_*ν*_:

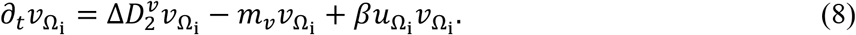

#### 2.2.2 Pest dynamics

We consider that edges do not modify directly pest population dynamics. Writing 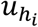 for the pest density in an edge *h*_*i*_:

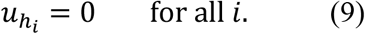

The pest is a specialist of the principal culture and, without dispersal, it shows positive growth only in crop fields. The bidimensional reaction-diffusion model for the pest density 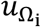 in field 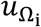 is

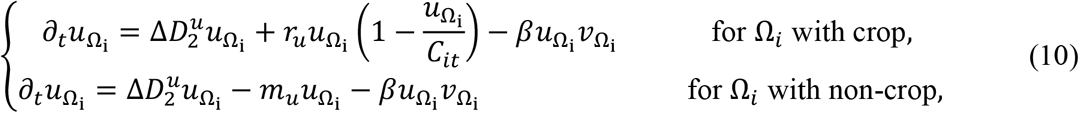

where 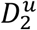 is the diffusion parameter of the pest in fields, *r*_*u*_ is its intrinsic growth rate on crop category, *β* is the predation rate, and *m*_*u*_ is the mortality rate of the pest on non-crop fields.

In a crop field, a pesticide treatment is applied when the average pest population density in that field exceeds a given threshold, which we here fix to 0.2 individuals/m^2^ per small unit of space and time. Pesticide treatments reduce the carrying capacity *C*_*it*_ of the field *i* (equation (10)):

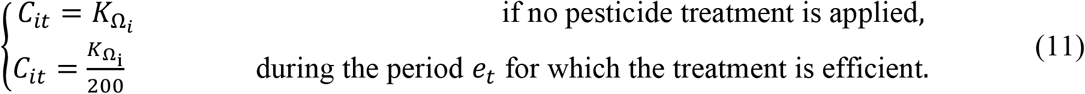

An additional mortality term could be added to account for the effects of pesticide treatments, but it would have implied the modification of both growth and carrying capacity. For that reason and to keep the model parsimonious, possible effects of pesticide treatments are assumed to change only the carrying capacity (Eq. 11). Similarly, mortality other than for predation or pesticide treatments could have occurred in crop fields, but we have opted against this option for the sake of parsimony.

#### 2.2.3 Coupling predator-pest dynamics over the entire landscape

Using the framework described in (11), the dynamics described by equations (7) to (10) are coupled over the full landscape using the following assumptions (see the Supplementary Information (SI) for more details): (i) edges (with or without a hedge) do not represent a barrier for the pest, (ii) edges without a hedge do not represent a barrier for the predator, (iii) the predator is attracted by hedges, thus migration from fields to hedges (*ρ*_21_) is relatively high, (iv) the predator shows aversion to move outside its natural habitat, thus migration from hedges to fields (*ρ*_12_) is lower than migration from fields to hedges. We consider reflecting conditions on landscape boundaries, meaning that in- and out-fluxes between the landscape and its surrounding environment are equal.

#### Inoculation and spatio-temporal design

Initially, the predator is present in all hedges at carrying capacity. The pest is introduced randomly in space and time. The average number of pest inoculations in a single simulation is proportional to the proportion of crop field area in the landscape, and we draw the actual number of inoculations from a Poisson distribution. The maximal average number of pest inoculations was fixed to 25 when the crop is grown in all fields. Inoculated crop fields are picked at random with probability depending on their relative surface.

### 2.3 Statistical methods for analysing simulation outputs

We define an experimental design based on Sobol’s sequences leading to 11,500 distinct parameter configurations. For each parameter combination, we consider 15 landscape replicates, leading to a total of 172,500 simulations.

We first conduct a Sobol sensitivity analysis on the mean and standard deviation of predator density, pest density and number of treatments by averaging the outputs over landscape replicates and crop fields. First-order indices were estimated with Sobol–Saltelli’s method (28,29), whereas total indices are estimated with Sobol–Jansen’s method (28,30). These analysis are performed within the R software v. 3.0.3 (R Team, 2003), using the packages fOptions (v. 3010.83) and sensitivity (v. 1.11).

Then, to further explore direction and magnitude of variations in response variables with respect to landscape parameters, we applied Generalized linear models (GLMs). Pest and predator densities, and pesticide treatment numbers (if different from 0), are analysed as response variable by using the Gamma distribution with log-link function. Additionally, pesticide treatment presence/absence during a simulation is analysed using a GLM with binomial distribution. We develop GLM formulas containing covariable interactions (see Table 1) up to 2nd order, and we use a step-wise variable selection algorithm based on the Bayesian information criterion (BIC) in order to select the “best subset” of variables for each model.

Finally, we use Generalized Linear Mixed-Effect models to analyse treatment occurrences by taking into account their spatial position in the landscape. We use the log-transformed area (*Area*) and perimeter (*Perimeter*) to take into account the geometrical properties of the fields, and we use the number of adjacent crop fields (*Adj*_*C*_), the number of adjacent hedges (*Adj*_*H*_), and the number of treatments applied in the adjacent crop fields (*Adj*_*Tr*_) to take into account the composition and dynamics in local neighbourhoods. In addition, we include the estimated linear effects from the global models as offsets. The random effect is structured by the landscape simulation to account for its specific dynamics. By analogy with the global GLMs, the presence/absence of treatments is analysed using the binomial response distribution, and pesticide treatment numbers are analysed with the Gamma distribution for the response variable with a log-link function. We consider predictor interactions up to 2^nd^ order. These analyses are performed using the R package lme4 with R version 3.2.3 (31).

## 3 Results

### 3.1 Sensitivity of predator density, pest density and pesticide treatments to model parameters

The sensitivity analysis of the mean of model outputs across landscape replicates (Figure 2a right) shows that variations in predator population density are mainly explained by predator migration 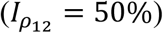 and by the proportion of hedges 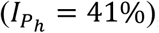, whereas interactions among parameters have little impact on the outputs. For the pest population density and the average number of pesticide treatments, crop proportion (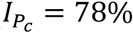 and 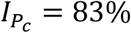, respectively) and pest growth rate (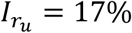 and 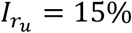, respectively) are the most important parameters to explain model output variability, again with only little interaction between model parameters (Figure 2b right). Complete results of pesticide treatments are in the SI.

**Figure 2:**
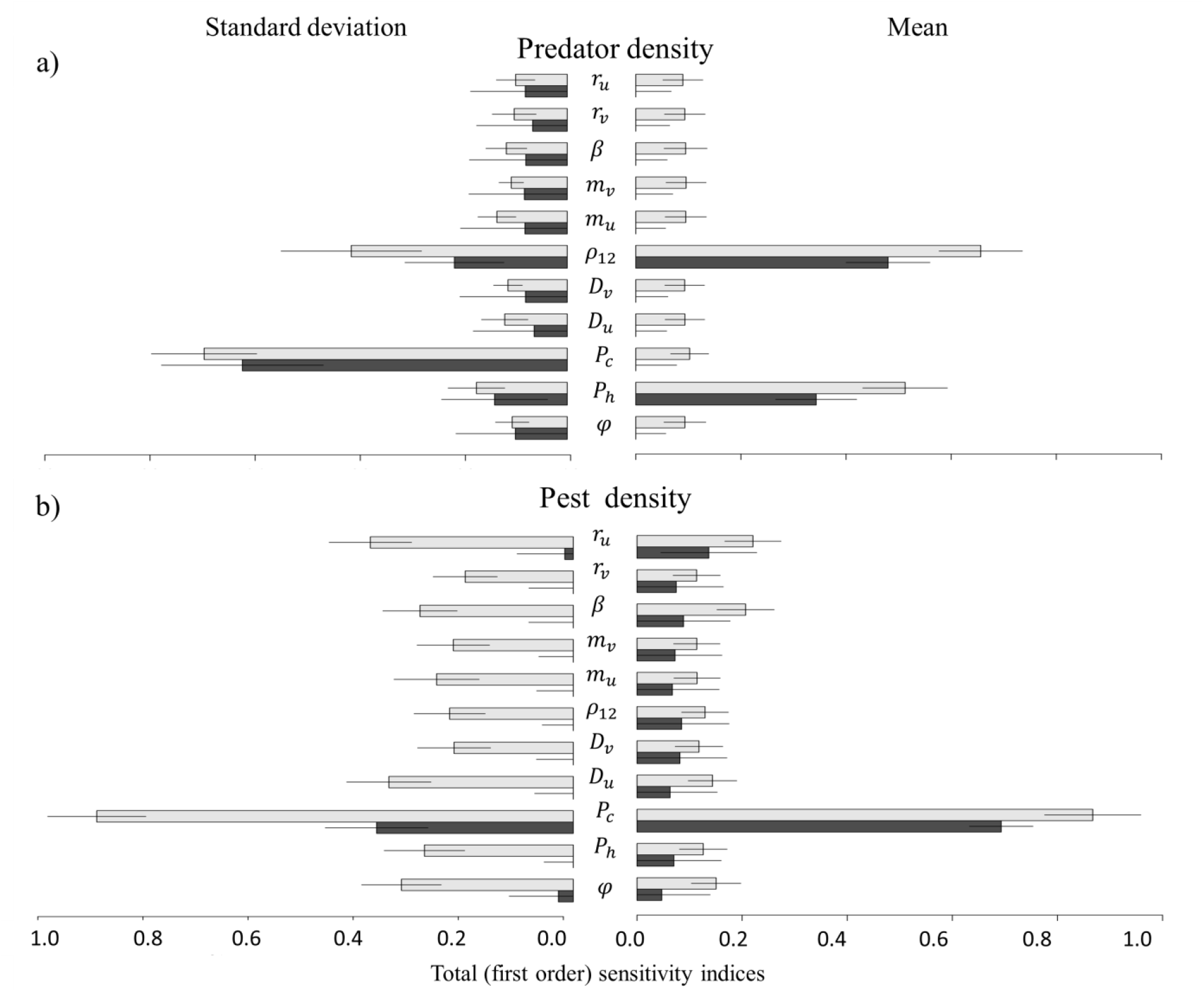
Sobol sensitivity analysis: Total sensitivity indices (light grey bar) and first-order sensitivity indices (black bar) of space-time averaged values for predator density (a) and pest density (b), based on the mean (right) or on the standard deviation (left) over replicated simulations. The length of the bar indicates the mean of the sensitivity index, and the solid line indicates its 95% confidence interval.

The sensitivity analysis of standard deviation of model outputs across landscape replicates gives different importance to the input variables as compared to the mean values. For the predator, crop proportion (*P*_*c*_), predator migration (*ρ*_12_), hedge proportion (*P*_*h*_) and spatial aggregation (*φ*) explain respectively 55%, 19%, 9% and 9% of the variability of model outputs (Figure 2a left). For the pest and pesticide treatments, results are consistent with the analysis on the mean. However, interactions between model parameters are important to explain variations of predator and pest density, as well as of pesticide treatments among landscape replicates. This implies that particular landscape structures characterized by a combination of several descriptors have to be considered to fully understand the drivers of predator-pest dynamics.

### 3.2 Landscape structure effects on the predator-pest dynamics

Effects of landscape variables (denoted *E*_*variable*_ in the following) on predator density highlight a positive effect of hedge proportion 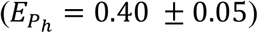, a negative effect of crop proportion 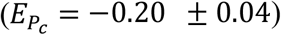 and a positive interaction among both variables 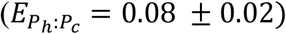, which implies that hedges can buffer the negative effect of increased crop proportion. Migration from hedges to fields 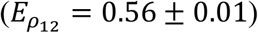 has the highest positive effect on predator density with again a positive interaction with crop proportion.

As expected, crop proportion 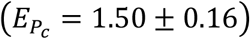, as well as spatial aggregation *(E*_*φ*_ = 0.55%0.02), have a strong positive effect on pest density. Both variables interact negatively 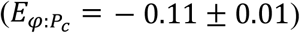, as high aggregation results in an increase of the size of contiguous crop fields, which lowers the effect of increased crop proportion. The positive effect of crop proportion is lowered by its interaction with hedge proportion 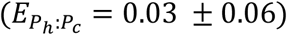 and also with predator migration from hedge to fields 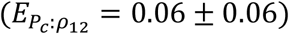. Counterintuitively at first sight, an increase in hedge proportion 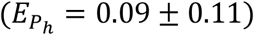 has a positive effect on pest population. Indeed, predator presence over all the landscape helps to stabilize the pest population by keeping it under the thresholds triggering a pesticide treatment. This is further confirmed by the fact that hedge proportion 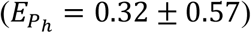, predator spillover 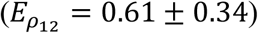 and concurrence of high crop proportion and aggregation 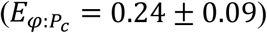 have a positive effect on the presence of pesticide treatments, but a negative effect on treatment numbers 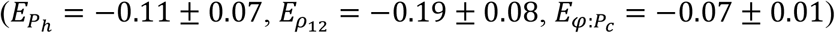.

Among species traits, predator migration from hedges to fields 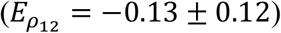 has the highest negative impact on pest density. Pest diffusion 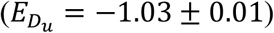, due to a dilution effect, and the predating rate (*E*_*β*_ = −0.24 ± 0.01), have also a negative impact on the pest, while the growth rate 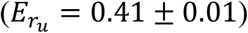 contributes positively to pest density. Figure S3 shows all estimated effects and confidence intervals for predator and pest density and treatment presence/absence and number, see also Table 2 for a sum up.

**Table 2:**
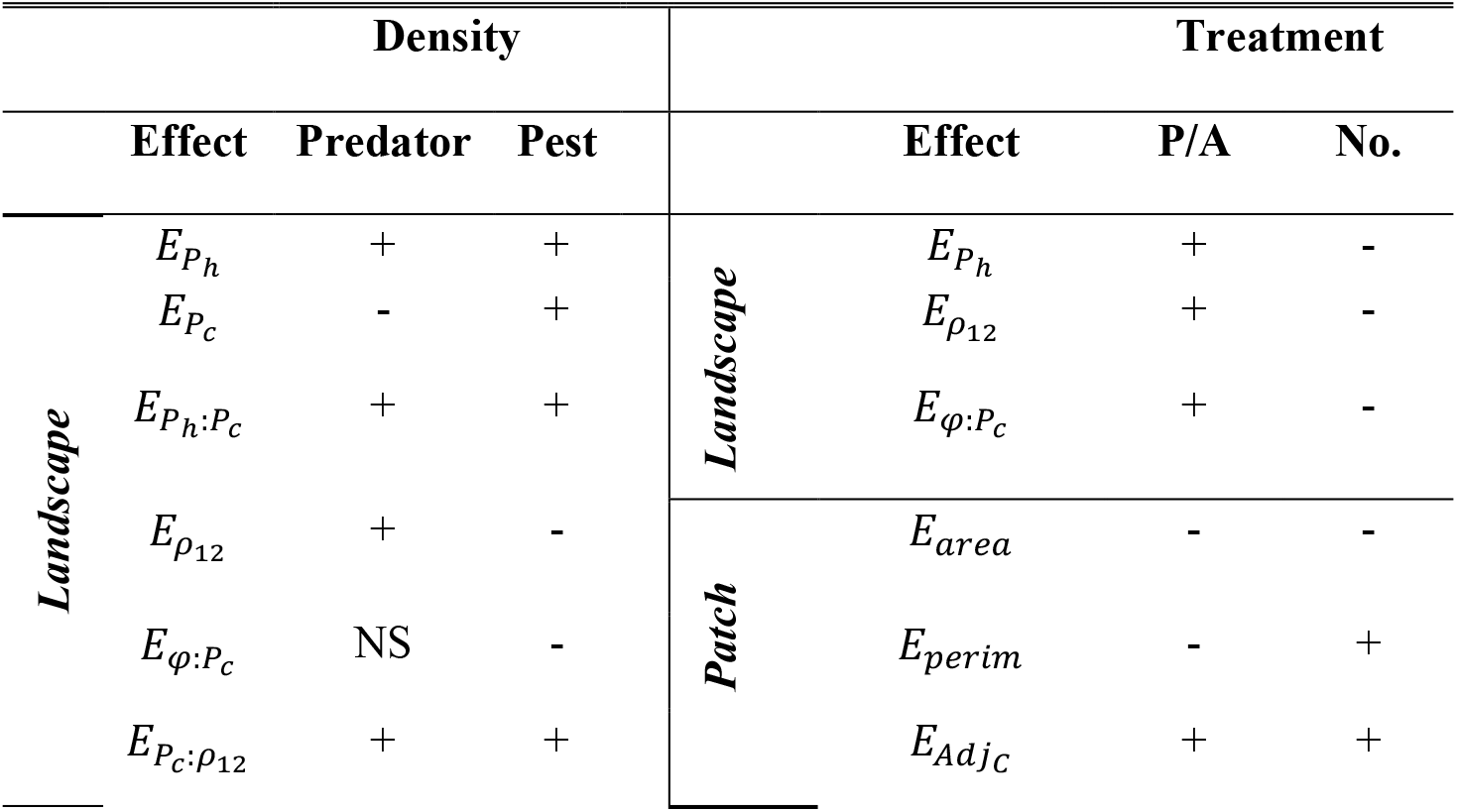
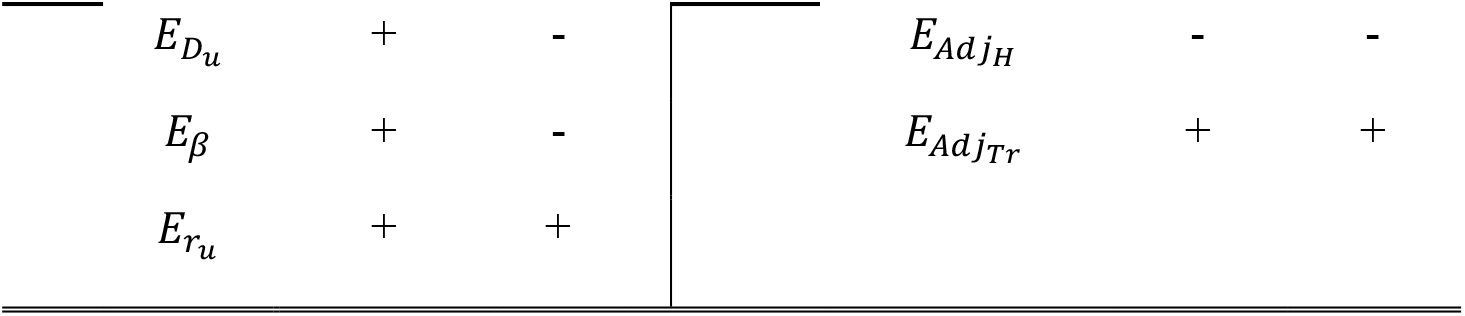
Summary of the presented estimated effects for predator and pest density (left) and for the presence/absence (P/A) and number (No.) of treatment (right) at landscape and field scale. + means positive effect, - means negative effect, NS means non significative effect.

| Density |  |  |  | Treatment |  |  |  |
| --- | --- | --- | --- | --- | --- | --- | --- |
|  | Effect | Predator | Pest |  | Effect | P/A | No. |
| Landscape | $E_{P_h}$ | + | + | Landscape | $E_{P_h}$ | + | - |
| | $E_{P_c}$ | - | + | | $E_{\rho_{12}}$ | + | - |
| | $E_{P_h:P_c}$ | + | + | | $E_{\varphi:P_c}$ | + | - |
| | $E_{\rho_{12}}$ | + | - | Patch | $E_{area}$ | - | - |
| | $E_{\varphi:P_c}$ | NS | - | | $E_{perim}$ | - | + |
| | $E_{P_c:\rho_{12}}$ | + | + | | $E_{Adj_C}$ | + | + |

|  |  |  |  |  |  |
| --- | --- | --- | --- | --- | --- |
| $E_{D_u}$ | + | - | $E_{Adj_H}$ | - | - |
| $E_{\beta}$ | + | - | $E_{Adj_{Tr}}$ | + | + |
| $E_{r_u}$ | + | + | | | |

### 3.4 Effect of local landscape on pesticide treatment

Presence of pesticide treatments is negatively influenced by field area and perimeter (*E*_*Area*_ = −0.32 ± 0.01, *E*_*Perimeter*_ = −0.10 ± 0.03). These effects reflect both a slower pest diffusion in large fields and a higher spillover of predators in fields with long perimeter. Conversely, when treatments occurred in a field, their total number increases with field perimeter due to spillover form the neighbourhoods. An increase in the number of adjacent crop fields produces a positive effect on the presence 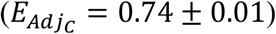 and number 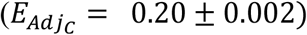 of treatments while an increase in the number of adjacent hedges leads to a negative effect on the presence 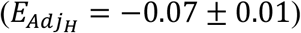 and number 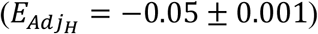 of treatments. Whereas in the global model the increase of hedge proportion is associated with a positive effect on the presence of treatments, we attribute the negative effect at local level to the fact that the predator tends to locally maintain the pest density under the application threshold, especially after a first treatment. The number of treatments in adjacent fields is positively correlated to the local presence 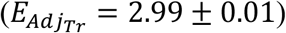 and number 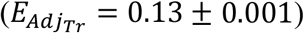 of treatments, indicating local pullulation of the pest. Figure S4 shows all estimated local effects and confidence intervals for treatment presence/absence and number, see also Table 2 for a sum up.

## 4 Discussion

Sustainable management of pests and diseases in agro-ecosystems requires a better understanding of how landscape structure drives and alters population dynamics. By simulating different landscape configurations including linear corridors, and the predator-pest dynamics, the present research aims at characterizing the joint influence of landscape structure and species traits on CBC service. Our study corroborates that spatial heterogeneity, landscape structure (i.e., the size and physical arrangement of patches), species traits and their interactions play a key role for CBC.

Crop proportion is the major determinant of increasing pest population and resulting in an increased number of pesticide treatments over the whole landscape. Indeed, increasing crop proportion in fragmented landscapes ensures food availability to the pest all over the landscape (1,2,12). In highly aggregated landscapes, the size of contiguous crop patches is already large enough to sustain a relatively large pest population, thus lowering the effect of an increase in crop proportion (14). The effects of crop proportion and spatial aggregation are intimately linked to pest growth rate and dispersal capability. Indeed, unfavourable landscape properties for the pest (i.e., low proportion and high fragmentation) can be compensated by a higher growth rate. However, the effect of dispersal is a double-edged sword since high dispersal helps spreading on fragmented landscapes but comes with a larger amount of propagules lost in unsuitable habitats, potentially leading to a dilution effect (3,33,34). As expected, hedge proportion (i.e., SNHs) positively affects predator presence in agricultural landscapes. In addition, the predator’s ability to move between SNHs and crop habitats is the parameter that increases most strongly the predator density, since it enables predators to reach complementary resources in crop fields more easily. Spillover from adjacent habitat is reported to have a major impact on pest populations in crop fields (3,12,34). Spillover not only depends on predator propensity to forage outside their natural habitat, but also on semi-natural patch connectivity and on crops and natural enemy reservoir interface (35). Thus, different combinations of SNH proportion and aggregation influence landscape structural connectivity and are also important determinants of predator efficiency in regulating crop pests (35).

In our representation, hedges are modeled as source of predators where they have a logistic growth. This is a simplification for predator dynamics in their natural habitat, as we do not consider potential prey presence in hedges and predator foraging behaviour in crop fields. For example, the growth rate, instead of being constant, could depend on the time spent in the fields and on the number of consumed preys. In addition, predating rate and consumption rate are crucial in determining the efficiency of CBC (36). Here, these parameters are not identified as influent in the dynamics, perhaps because they are assumed identical (parameter *β* in our model). Finally, another strong assumption may be that pesticide treatments do not affect predator mortality. However, in crop fields a positive predator growth rate relies only on pest, such that a strong pest reduction due to treatments is automatically translated into a strong impact on predator density when such treatments occur. Thus, adding extra-mortality in crop fields should not modify the results that much. To account for major pesticide treatment effects, an impact on predator mortality in its natural habitat should be considered.

In our analysis, we found that predator ability to disperse from hedges to crop fields has a major influence on pest density and related treatments. High crop proportion enhances pest density, but this effect is counter-balanced by the joint effect of hedge proportion and predator spillover, which favours predator pressure and reduces pesticide treatment application. Indeed, hedges ensure an increased landscape functional connectivity, which enables predators to successfully disperse and feed on complementary resources in the fields. Interestingly, however, we found that if SNHs can sustain a high population of natural enemies (37), this is not sufficient to achieve a decrease in pest density. Indeed, by keeping the pest population density under the treatment threshold, the predator population can favour its spread across the landscape, thus increasing pest density at the landscape scale, even if fewer treatments are applied. Most of the studies consider the amount of SNH as a proxy for predator presence and focus on how landscape structure directly influences CBC. However, as highlighted by our results (see also 37), the extent to which species are influenced by landscape heterogeneity depends on their traits. For example, (39) argue that natural enemies with an oriented movement are better able to deliver pest control services. They discuss the interplay among predator mobility, proportion of crop and SNHs. More generally, SNH predator spillover is expected to be particularly strong when (i) predator attack rates on prey are high, (ii) predator movement abilities are substantial, and (iii) predator mortality rates in the recipient habitat are low (40).

The amount of predator spillover, and the distance over which pest and predator can spread, both depend on local configurational variables such as field size, shape, amount of shared edge, and connectivity (19). Large fields can support high pest volumes, but it has been demonstrated that the relationship between field size and pest density can take several forms depending on assumptions, conditions and species considered (41). Our results show a negative effect of large field area on the need to use pesticides and on the number of required applications, which, accordingly to (41), may come from the elevated growth rate of the prey combined with its good dispersal ability. By contrast, the perimeter shows a positive relationship with pest density by increasing treatment numbers due to the spillover from surrounding fields. However, when long perimeters are coupled with hedge presence on them, predator spillover into fields reduces pesticide treatment numbers (9). Interestingly, we show a contrasted effect of hedge depending on the scale considered. At global scale, the proportion of hedges shows a positive effect on pest density and has a negative effect only on treatment application presence. At local scale, the number of edges adjacent to a crop field shows an even more important impact on CBC by negatively affecting both the local presence and number of treatments (37).

Landscape simplification is a major driver of pest abundance and consequently has strong impacts on the necessity of pesticide treatments and their frequency. We find that natural habitat enhances predator population, but it does not systematically translate into a strong correlation with pest density decrease. However, a relatively high predator density often helps maintaining pest density below the economic threshold level above which pesticides are applied, thus preventing highly localized pest densities. By contrast, predator spillover from hedges to fields is fundamental for CBC; it reduces pest density and guarantees high predator fluxes and different habitat connectivity. At field scale, landscape geometrical features, hedge presence and habitat connectivity are able to influence predator-pest dynamics, and therefore they affect the number of pesticide treatments. This highlights the importance of conducting a multi-scale analysis to consider the differences in outcomes at landscape and patch scale for pest CBC (14). In most of our analyses, we considered global outputs by averaging pest and predator densities over crop fields. However, populations are obviously structured in space and time. Thus, a complementary analysis studying how landscape structure impacts predator-pest spatio-temporal dynamics would bring more insights on pest outbreak determinants. Moreover, a larger number of pest and predator species, inter/intra-species interactions and also different trophic network structures, could be considered in future work to better understand the role of pest and predator diversity on CBC efficacy.

## Supporting information

Supplementary material: model details and further results

## Acknowledgments

We are thankful to Claire Lavigne and Katarzyna Adamczyk for their help and wise suggestions for landscape data processing and for the discussion part. Khadija Sabir has been supported by the Erasmus+ - KA1 Erasmus Mundus Joint Master Degrees Programme of the European Commission under the PLANT HEALTH Project.

