## Supplementary material: model details and further results for "More pests but less treatments: ambivalent effect of landscape complexity on Conservation Biological Control"

### Supplementary Information (SI) of the paper: "Understanding the effects of complex agricultural landscapes on conservation biological control: A stochastic-mechanistic dynamic modeling approach"

#### SI1. Description of the 2D/1D model for population dynamics in the landscape

Here, we detail the description of the dynamics of a species in a landscape defined as a 2D matrix crossed by 1D corridors following the methodology developed in Roques & Bonnefon, 2016. Here, we report some of the key information to understand the 2D/1D model for population dynamics for our analysis. More details are reported in the original paper Roques & Bonnefon, 2016.

2D reaction-diffusion equations describes the dynamics in the matrix, another set of 1D reaction-diffusion equations describes the dynamics in the corridors. The fluxes among the matrix and the corridors are described by coupling terms between the two sets of equations.

We consider a 2D matrix defined by a set $\Omega \subset R^2$ , composed by finite mosaic $i$ of polygonal disjoint 2D patches $\Omega_i$ separated by corridors (Figure 1). Patch boundary is denoted by $\delta\Omega_i$, each boundary consisting of a finite number of 1D edges $\lambda_i^k$ . The edges can be classified as: the interior edges (= the corridors), and the exterior edges which belong to the boundary $\delta\Omega$ of $\Omega$ and where no particular 1D dynamics is modelled. The population density is denoted by $v_i$ in each patch $\Omega_i$ and by $u_i^k$ in each corridor $\lambda_i^k$.

.

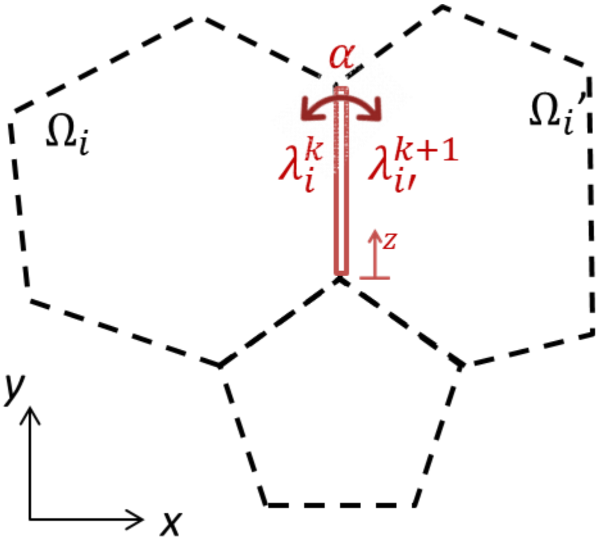


Figure 1: Landscape representation defined by patches $\boldsymbol{\Omega_i}$and edges $\boldsymbol{\lambda_i^k}$over patch boubdary.

##### 1.1 Dynamic in the matrix

The population density is modelled by a reaction-diffusion equation:

$$\delta_t v_i = d\Delta v_i + f(v_i)$$

$d$ is the diffusion parameter that describes the mobility in the matrix 2D, $f$ is the growth function that describe the birth and death events in the patch $\Omega_i$.

The exchange among patch $\Omega_i$ and the surrounding corridors are described by the fluxes terms:

$$d\nabla v_i\,\boldsymbol{n} = \rho_{12}u_i^k(t,x,y) - \rho_{21}v_i\ (t,x,y)$$

$\rho_{12}u_i^k(t,x,y)$ describes the flux of individuals leaving the corridor $\boldsymbol{\lambda_i^k}$ and entering the patch $\boldsymbol{\Omega_i}$ at time $t$ and at the position $(x,y)$, $\rho_{21}v_i\ (t,x,y)$ describes the flux of individuals leaving the patch $\boldsymbol{\Omega_i}$ and entering the corridor $\boldsymbol{\lambda_i^k}$, $\boldsymbol{n} = \boldsymbol{n}(\boldsymbol{x},\boldsymbol{y})$ denotes the outward unit normal to the boundary $\delta\Omega_i$. On the exterior boundary edges $\boldsymbol{\lambda_i^k} \in \delta\Omega_i$ standard reflecting boundary conditions are assumed: $d\nabla v_i\,\boldsymbol{n} = 0$. These boundary conditions mean that either the individuals crossing the boundaries are reflected inside the domain.

#### 1.2 Dynamics in the corridors

Each corridor $\boldsymbol{\lambda_i^k}$ belongs to the common boundary of $\Omega_i$ and of another set, which is denoted by $\Omega_{i'}$, *i.e.*, $\boldsymbol{\lambda_i^k} = \boldsymbol{\lambda_{i'}^{k'}}$, in way to model the 1D dynamics on each side of the corridor. The population densities in the corridor can be denoted by $u_i^k$ and $u_{i'}^{k'}$ from the $\Omega_i$ side and the $\Omega_{i'}$ side respectively, and we assumed that $u_i^k \neq u_{i'}^{k'}$, in general. The exchanges between the two sides of the corridor are taken into account through a permeability parameter $\alpha > 0$ (Fig. 1). To state the 1D equation for the dynamics in the corridors, we define an isometric transformation $z \to \big(x(z), y(z)\big)$ which maps any corridor $\boldsymbol{\lambda}$ into an interval $(0, L(\boldsymbol{\lambda}))$, where $L(\boldsymbol{\lambda})$ is the length of the corridor. Thus, the population density in the new coordinate $z \in L(\lambda)$ is defined by $\tilde{u}(t,z) = u(t,x,y)$. The population dynamics in each corridor $\boldsymbol{\lambda_i^k} = \boldsymbol{\lambda_{i'}^{k'}}$ separating two patches $\Omega_i$ and $\Omega_{i'}$ is given by:

$$\delta_t\tilde{u}_i^k = D\delta_{zz}\tilde{u}_i^k + \rho_{21}v_i\ \big(t, x(z), y(z)\big) - \rho_{12}\tilde{u}_i^k(t,z) - \alpha\tilde{u}_i^k(t,z) + \alpha\tilde{u}_{i'}^{k'}(t,z) + g\big(\tilde{u}_i^k\big),$$

$$t > 0, \qquad z \in (0, L(\boldsymbol{\lambda_i^k}))$$

$g$ is the growth function in the corridor $\boldsymbol{\lambda_i^k}$; $\alpha$ permeability parameter among the two side of the corridors, D is the diffusion parameter on 1D corridor.

**SI2. Complete Sobol sensitivity analysis for predator and pest density and pesticide treatment.**

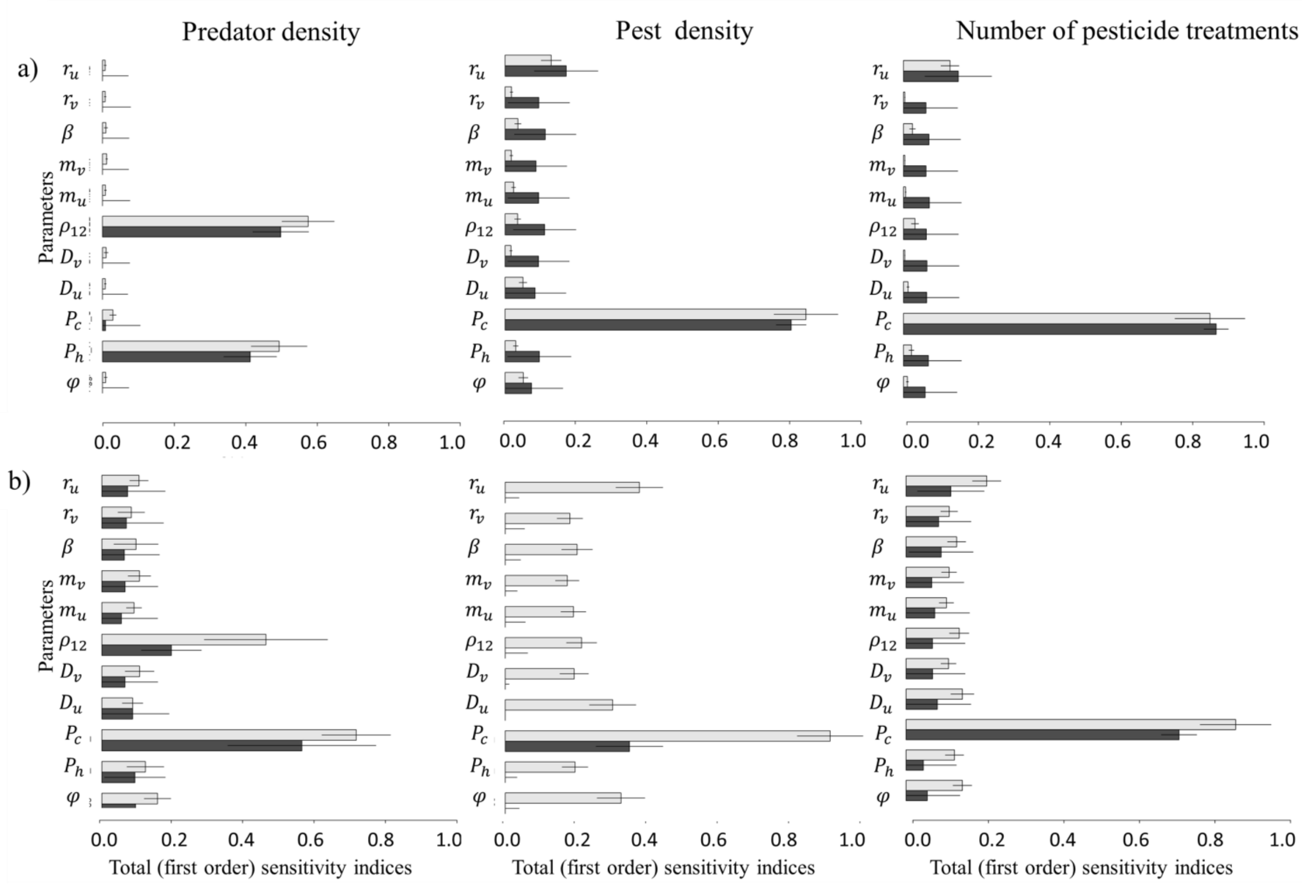


Figure 2: Sobol sensitivity analysis: Total sensitivity indices (grey bar) and first-order sensitivity indices (black bar) of space-time averaged values for predator density, pest density and number of pesticide treatments based on the mean (Panel a) or on the variance (Panel b) over replicated simulations. The length of the bar indicates the mean of the sensitivity index, and the solid line indicates its 95% confidence interval.

**SI3. Estimated effects of Generalized linear models (GLMs) for pest and predator densities, and pesticide treatment presence/absence and number and Generalized Linear Mixed-Effect for local pesticide treatment presence/absence and number.**

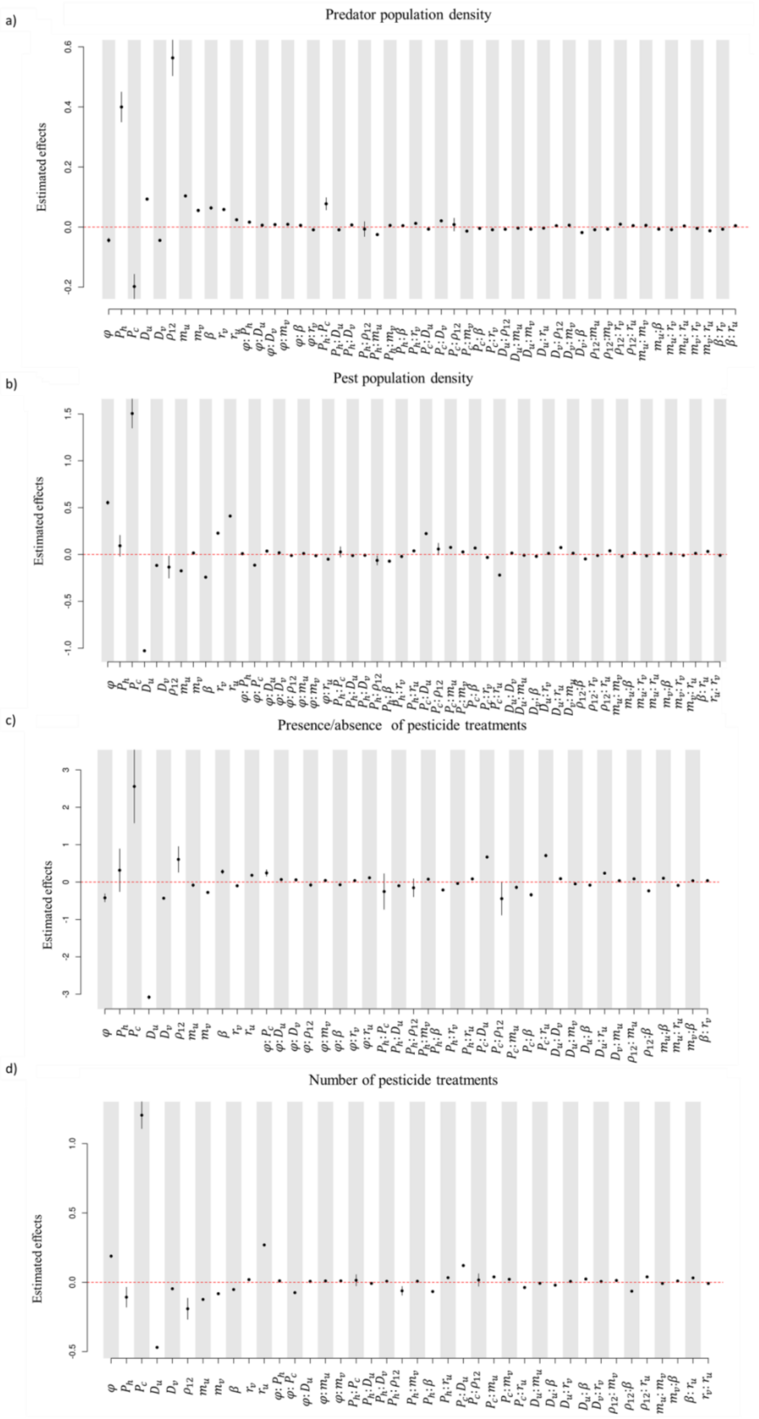


**Figure 3: GLM coefficient estimates.** Effects of input parameters and their bivariate interactions on pest and predator population dynamics: Coefficient estimates (dots) and their confidence intervals (segments) for the parameters retained by the stepwise selection in the GLM for the predator density (a), the pest density (b), the presence/absence of pesticide treatments (c) and the number of pesticide treatments (d).

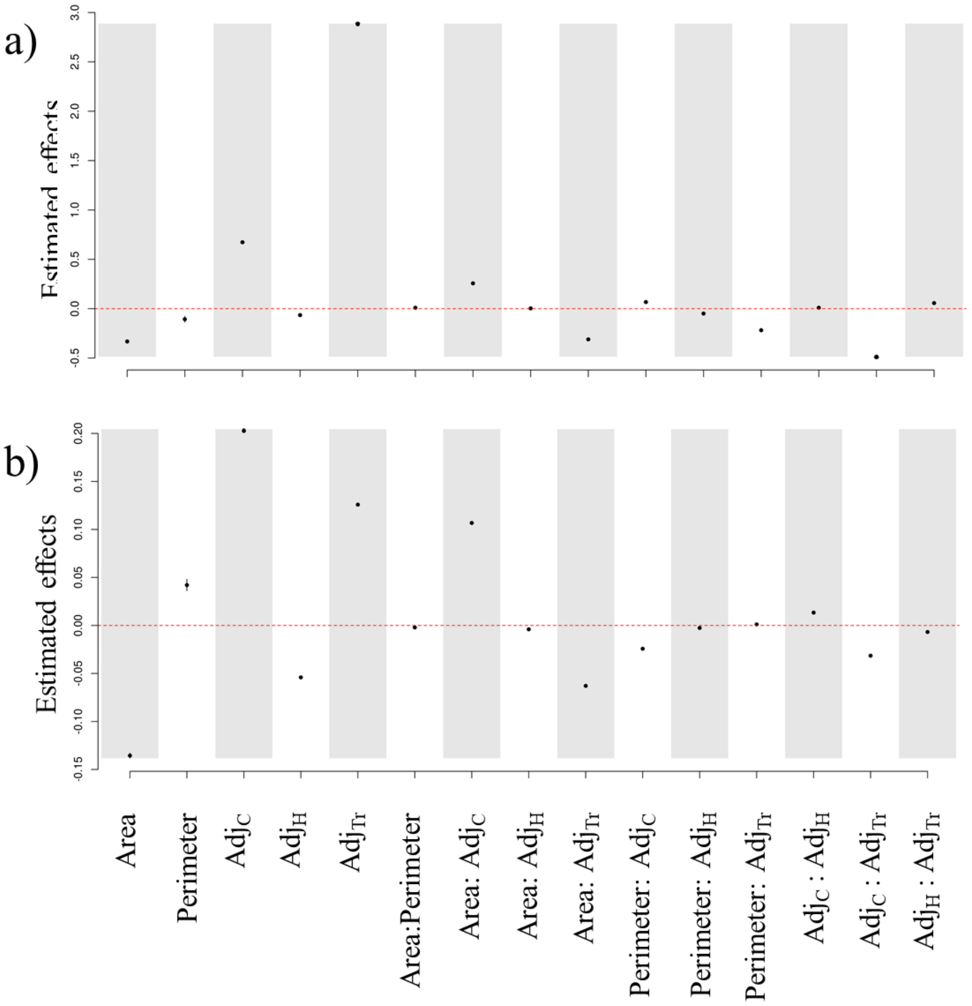


**Figure 4: Generalized Linear Mixed-Effect coefficient estimates.** Estimated local effects (dots) and confidence intervals (segments) for the presence/absence of treatments (a) and for the number of treatments (b). The intercept values are not shown in this plot to better focus on the effects of the landscape covariables.
